# A novel approach to the detection of unusual mitochondrial protein change suggests low basal metabolism of ancestral anthropoids

**DOI:** 10.1101/2021.03.10.434614

**Authors:** Bala Anı Akpınar, Vivek Sharma, Cory D. Dunn

## Abstract

The mitochondrial genome encodes core subunits involved in the process of oxidative phosphorylation. The sequence and structure of these mitochondria-encoded polypeptides are expected to be shaped by bioenergetic requirements linked to diet and environment. Here, we have developed a robust and effective method for highlighting phylogenetic tree edges with unexpectedly rapid, and likely consequential, substitutions within mitochondrial proteins. Further, our approach allows detection of discrete protein substitutions likely to alter enzyme performance. A survey of mammalian taxonomic groups performed using our method indicates that widely conserved residues in mitochondria-encoded proteins are more likely to rapidly mutate toward variants providing lower OXPHOS activity within specific clades. Intriguingly, our data suggest reduced cellular metabolism of ancestral anthropoids, and our findings have potential implications regarding primate encephalization.

**Significance Statement:** Mitochondria harbor DNA (mtDNA) that encodes proteins important for converting food into energy. The environment and lifestyle of an organism shapes, and is shaped by, the sequences of these mitochondrial genomes. We developed a new approach for the detection of rapid functional change to proteins, and we applied our method to the mitochondria-encoded polypeptides of mammals. We found that primates displayed a general signature of relative hypometabolism that is shared with other mammals characterized by a low metabolic rate. Indications of reduced cellular metabolism extend even to the earliest anthropoids. Our findings have potential implications regarding the evolution of an enlarged primate brain.

## Introduction

Mitochondria are derived from an ancestral endosymbiont related to alpha-proteobacteria (Bonen et al. 1977; Roger et al. 2017). While most of the genes encoded by the pre-mitochondrial endosymbiont were lost or transferred to the (proto-)nucleus, a handful of protein-coding genes continue to reside upon a mitochondrial genome that is inherited maternally in vertebrates (Hoekstra 2000; Christie and Beekman 2017). The mitochondrial DNA (mtDNA) of mammals typically encodes 22 transfer RNAs, 2 ribosomal RNAs, and 13 proteins that are crucial for oxidative phosphorylation (OXPHOS) (Gustafsson et al. 2016; Pfanner et al. 2019). During the process of OXPHOS, energy derived from food drives a proton gradient across the mitochondrial inner membrane in a series of reactions leading to oxygen consumption (Sousa et al. 2018). This proton gradient is then harnessed by a mechanoenzyme to regenerate ATP.

Intra- and inter-species comparisons of endotherm physiology most commonly focus upon the basal metabolic rate, or the measured oxygen consumption in an animal ‘at rest’ at thermoneutral conditions (Schmidt-Nielsen 1997; Konarzewski and Książek 2013). Frequently, basal rates of metabolism are converted to mass-specific rates reflecting oxygen consumption per unit of body mass (McNab 1999). As the vast majority of oxygen consumption occurs during OXPHOS (Larsen et al. 2011), basal metabolism should be determined, to a great extent, by the sequence and corresponding structure of mtDNA-encoded proteins. Contrast between a particular mammal’s basal metabolism and an expected value emerging from the strong allometric relationship between basal metabolic rates and body mass (Kleiber 1932; White and Seymour 2003) can often be predictive of other physiological and ecological factors, such as body temperature, mode of locomotion, endurance, running speed, diet, and other key aspects of life history and habitat (Lovegrove 2000, 2004; McNab 2012; Konarzewski and Książek 2013; Swanson et al. 2017). Consequently, these additional aspects of metabolism and physiology should also be expected to be determined by the mitochondrial genotype. Supporting this idea, changes in mitochondria-encoded polypeptides have been identified within particular clades that may be linked to their metabolic needs. For example, unexpected substitutions to mtDNA-encoded proteins potentially associated with transitions in environment or physiology have been identified within multiple vertebrates (Garvin et al. 2015), including bats (Shen et al. 2010), primates (Pierron et al. 2012), hummingbirds (Dunn et al. 2020), ‘high-performance’ fish (Dalziel et al. 2006), snakes (Castoe et al. 2008), and aquatic mammals (McClellan et al. 2005).

Here, we have developed a straightforward, empirical, and computationally undemanding method that can highlight phylogenetic tree edges and polypeptides characterized by greater-than-expected and potentially consequential change to conserved amino acid positions. We have applied our approach to the full set of mammalian reference mtDNAs at multiple taxonomic levels, and we found that specific clades tend to be enriched for mutations predicted to affect mitochondrial function. Primates clearly possess an elevated propensity for substitution at key locations within mitochondria-encoded proteins. Further analyses indicate that ancestral anthropoids are likely to have been characterized by low basal metabolic rates relative to many other mammals. Our results allow speculation regarding the evolutionary path that primates have taken toward greater encephalization.

## Results

Toward a comprehensive analysis of mitochondrial protein evolution in mammals, we recovered full GenBank records for mammalian mtDNAs stored in the RefSeq database (O’Leary et al. 2016). The mtDNA record for the reptile *Anolis punctatus* was also retrieved for the purpose of rooting a mammalian phylogenetic tree. Coding sequences were extracted and concatenated, a practice consistent with uniparental inheritance and co-evolution of vertebrate mtDNA-encoded polypeptides (Sato and Sato 2017). Concatenated protein-coding sequences were aligned using MAFFT (Katoh and Standley 2013), and phylogenetic tree inference was performed using RAxML-NG (Kozlov et al. 2019). Next, we used the PAGAN package (Löytynoja et al. 2012) to reconstruct the amino acid character values of each position at all bifurcating nodes within our maximum likelihood mtDNA tree. Output protein alignments from PAGAN were ungapped and indexed using *Bos taurus* protein sequences as a reference, a common approach prompted by early and extensive use of this species during structural and functional characterization of mammalian OXPHOS complexes. Using the tree and node character values emerging from PAGAN, we inferred all amino acid substitutions at positions for which less than 2% of extant and ancestral sequences contained gaps (Supplementary File 1). Taxonomic information for each input sequence was recovered from the National Center for Biotechnology Information (NCBI) (Schoch et al. 2020), and taxonomic information at internal nodes was assigned by concordant annotation of all descendent external tree nodes at each selected taxonomic level. The resulting taxonomic dataset was then linked to each analyzed tree edge and amino acid substitution.

Using our table of mitochondrial protein substitutions, we determined the relative conservation of each analyzed amino acid position by calculating a total substitution score (TSS, Figure 1A), which is simply the sum of amino acid substitution events mapped to a given alignment position across all tree edges. Importantly, the dynamic range of TSSs is potentially unlimited and can take advantage of the continuous accretion of new sequence information. This ‘tree-aware’ metric may surpass measures of conservation that consider positional character frequencies extracted from multiple sequence alignments, since character frequencies are sensitive to sampling biases. The TSS avoids taking explicit account of amino acid side chain chemistries, since the ability to replace one amino acid for another within a polypeptide can be highly context-dependent (Zuckerkandl and Pauling 1965; Dunham and Beltrao 2021). However, the number of substitutions at a site must implicitly reflect the underlying biochemical consequences of replacing one amino acid for another. The minimum possible TSS for an amino acid position is zero, if no substitutions have been detected at a given position, and the maximum possible TSS for an analyzed position would correspond to the total number of tree edges (here, 2492 total edges). When plotting the TSSs obtained during our analysis of mammalian mtDNA-encoded proteins, it is clear that the vast majority of these mitochondria-encoded amino acids are quite conserved and seldom change over roughly 200 million years of evolution within this vertebrate class (Figure 1B). Indeed, the median TSS of analyzed positions is 9, the TSS at the 25th percentile is 1, and the TSS at the 75th percentile is 56. However, a smaller subset of aligned, sparsely gapped positions was more tolerant of substitutions, and the maximum TSS obtained across all analyzed mtDNA-encoded amino acid positions was 361. TSSs for each mitochondrial protein position, calculated from the mammalian dataset and indexed using the *Bos taurus* reference sequence, are provided as Supplementary File 2.

**Figure 1:**
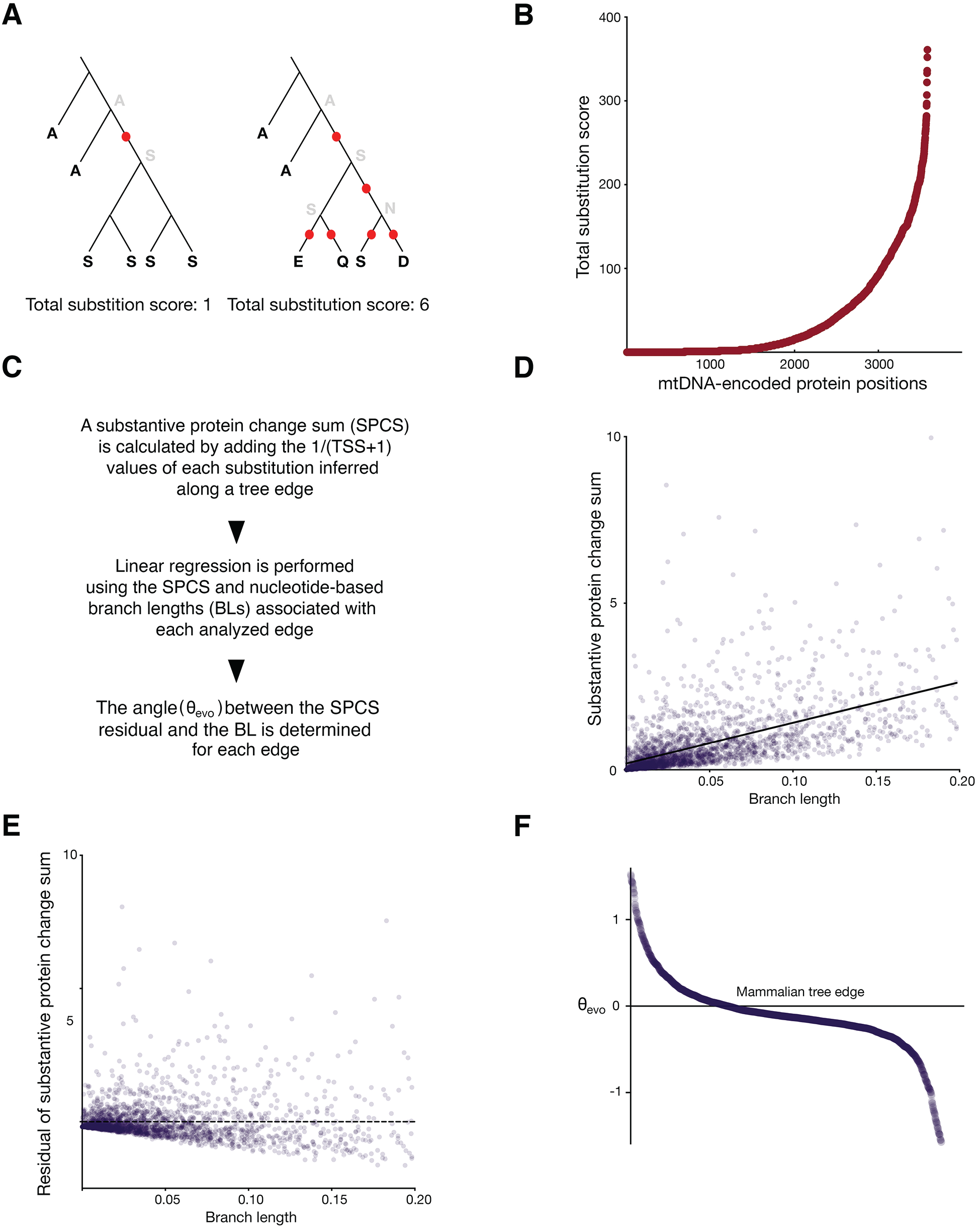
Detection of accelerated and potentially functional protein evolution along phylogenetic tree edges. (**A**) The total substitution score (TSS) is a measure of alignment position conservation. Character values at internal nodes are predicted, and inferred substitutions along an edge are summed. (**B**) The TSSs of all alignment positions gapped within less than 2% of inferred ancestral and input sequences are plotted for all mitochondria-encoded proteins. Alignment positions are ordered by increasing TSS. (**C**) The methodology for detecting whether an edge is characterized by greater-than-expected protein evolution is outlined: (**D**) The alignment position TSS for each substitution detected along an edge was added to 1, and the reciprocals of these values were summed to generate a substantive protein change sum (SPCS) for each edge found within a tree of mammalian mtDNA sequences. Edges with a branch length, greater than or equal to 0.2 were removed, and linear regression was performed (black line). (**E**) Residuals for each sample were calculated, and (**F**) the arctangent of the residual divided by the branch length (θevo), for each analyzed edge across all analyzed mitochondria-encoded amino acid positions is computed.

Next, we extended our analysis to highlight those specific tree edges where substantive change to mammalian mtDNA-encoded proteins had occurred. Substitutions at a position with a high TSS (low conservation) and those occurring at a position with a low TSS (high conservation) might be expected to have effects of differing magnitude, and so any approach highlighting edges characterized by unexpectedly rapid protein evolution at otherwise conserved sites should take into account both the number *and* the potential effect of substitutions. Consequently, we recovered the TSS for each substituted amino acid position along a tree edge, then added 1 and calculated the reciprocal value in order to augment the impact of changes at the most conserved sites during our search for the most impactful change to protein activity. We then added these values together to generate a substantive protein change sum (SPCS) (Figure 1C). A subset of tree edges was clearly associated with an elevated SPCS when considering all analyzed positions within mtDNA-encoded proteins.

The SPCS should be of great assistance in identifying tree edges harboring an unexpected magnitude of functional change to proteins. However, when inferring functional effects from sequence changes, one should also attempt to take into account potential compensatory changes occurring over time that might ‘cushion’, or render functionally neutral, the identified substitutions (Kondrashov et al. 2002; Starr and Thornton 2016). To pursue this line of reasoning, we compared the SPCS of an edge to the evolutionary distance, represented by a tree <u>b</u>ranch <u>l</u>ength (BL) calculated using nucleotide sequence divergence and reported during our PAGAN analysis. Only edges with BLs averaging less than 0.2 substitutions per site were included within this regression analysis (Figure 1D) while longer edges were discarded. When plotting SPCSs against BLs, we encountered a linear relationship (R-squared of 0.32, p-value <0.0001) between SPCS and BL, with most tree edges (69%) falling below the regression line. SPCS residual magnitude appeared to scale with BL (Figure 1E), and some edges were clearly characterized by a higher-than-expected SPCS. These results are consistent with the idea that longer divergence times generally provide a greater opportunity for compensatory mutations, but that actual functional change can indeed be localized to particular tree edges. So that we might more meaningfully contrast SPCSs calculated from edges of low BL with those calculated from edges of high BL, we then determined the arctangent (expressed in radians) of the SPCS residual divided by the BL. The resulting angle (θ_evo_) is a normalized metric suggestive of functionally relevant polypeptide change.

Clearly, a number of edges exhibit a substantially elevated value of θ_evo_ when considering all mtDNA-encoded amino acid positions analyzed together (Figure 1F), or when analyzing individual respiratory complexes (Supplementary Figure S1). Demonstrating the utility of our approach, we encountered edges leading to several domesticated species among those with the highest θ_evo_ scores calculated across all subunits. Indeed, edges leading to the species *Bos taurus* (cattle), *Sus scrofa* (pigs), and *Ovis aries* (sheep) were ranked first, seventh, and eighth of 2329 analyzed edges (Supplementary File 3). These findings likely correspond with adaptation to the needs of their human captors or with potential population bottlenecks encountered during domestication (Larson and Fuller 2014; Bortoluzzi et al. 2020). Edges leading to several vulnerable or endangered species (IUCN 2008a, b, 2016b, a) are also commonly found within the top-ten ranked edges with respect to θ_evo_ calculated across all mitochondria-encoded proteins, including *Gorilla gorilla* (western gorilla), *Ursus thibetanus mupinensis* (Asian black bear, Sichuan subspecies), *Budorcas taxicolor* (takin), and *Panthera tigris amoyensis* (South China tiger). These data are, once again, consistent with population bottlenecks that allow persistence of unusual and potentially deleterious substitutions (Ohta 1973). Unusual substitutions at polypeptide positions that are, with the exception of the indicated edge, totally conserved among mammals contribute greatly to the high rank of these edges (Supplementary File 1). Our full θ_evo_ dataset, as well as the matching inferred substitutions, should serve as a valuable resource for those studying the evolution of mitochondrial proteins within specific mammalian groups, as well as their ancestors and kin.

Our method appears to successfully highlight individual tree edges harboring unusual and potentially impactful substitutions to mtDNA that might raise or lower the activity of different OXPHOS complexes. Next, to better understand selection on mtDNA-encoded proteins within various mammalian taxonomic groups, we carried out an analysis to reveal which clades might harbor an unusual number of tree edges with high θ_evo_ values. Toward this goal, we first calculated the median θ_evo_ for mammalian orders that were assigned five or more edges. Although orders typically encompassed individual edges exhibiting a wide diversity of θ_evo_ values (Figure 2A), several orders were clearly characterized by a greater-than-expected median θ_evo_. The three orders with the highest median θ_evo_ values were Proboscidea (encompassing elephants and mammoths), Pilosa (sloths and anteaters), and Primates (including monkeys, apes, lemurs, and lorises), while the three orders with the lowest median θ_evo_ values were Perissodactyla (odd-toed ungulates), Artiodactyla (even-toed ungulates), and Lagomorpha (rabbits, hares, and pika). Lower boundaries of the 90% confidence interval of the median θ_evo_ (Figure 2B) further indicated that rapid and substantive protein evolution was occurring within the mitochondria of Primates and Pilosa, while the upper boundaries of the 90% confidence interval of the median θ_evo_ (Figure 2C) supported constrained evolution of mitochondrial proteins in Artiodactyla, Lagomorpha, and Perissodactyla. Similar outcomes emerged during analyses of individual OXPHOS complexes (Supplementary Figs. S2 and S3)

**Figure 2:**
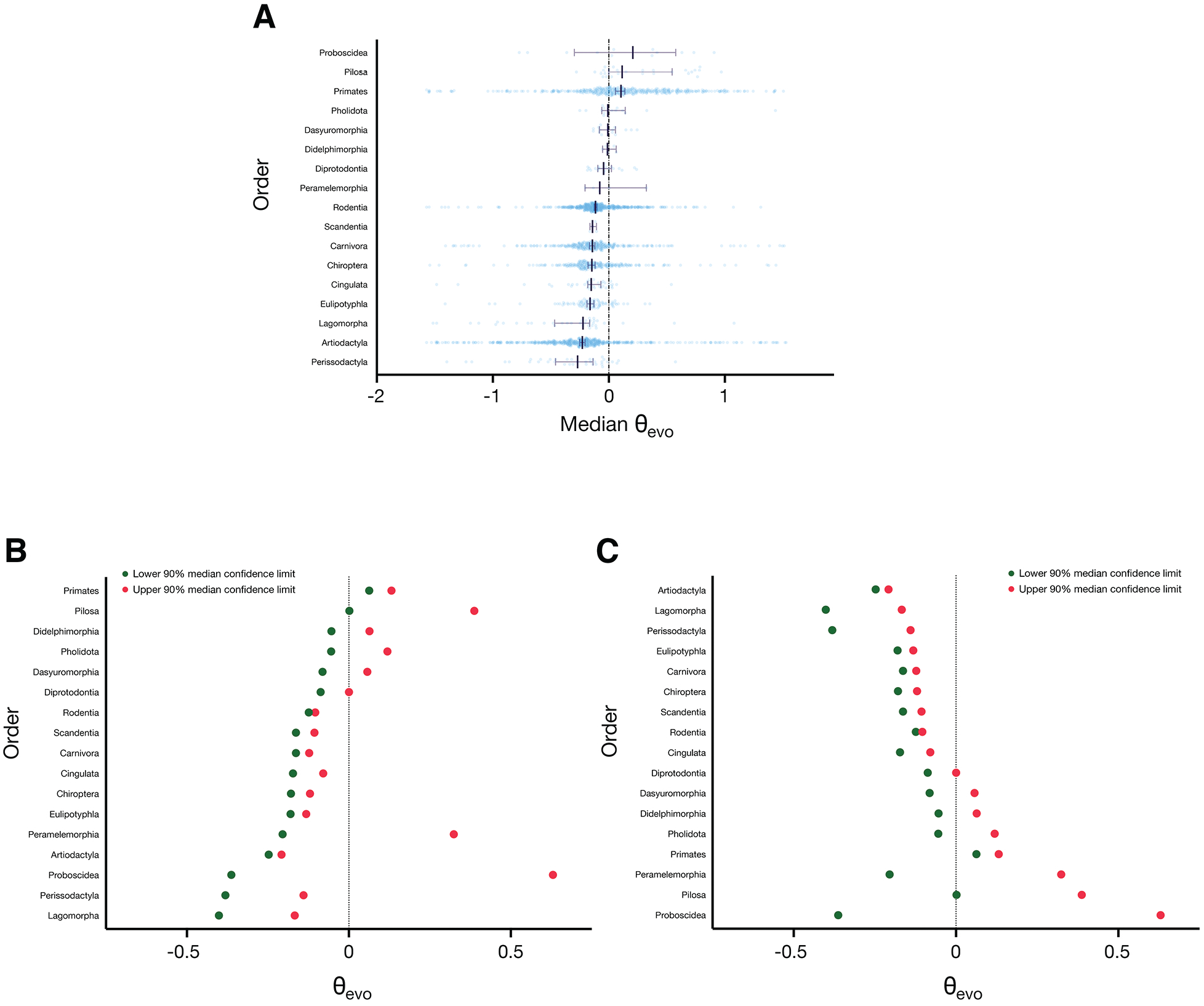
Mammalian orders differ in their propensity for potentially consequential mitochondrial protein substitutions. (**A**) The median θevo calculated for each order assigned five or more edges is shown (vertical bar), along with the interquartile range (error bars). Lower and upper 90% confidence intervals of the median are plotted, with the orders arranged from either the highest 90% median confidence limit to the lowest (**B**) or arranged from the lowest upper 90% median confidence interval to the highest (**C**).

Next, we directed our analyses toward mammalian families. As seen during our analysis of mammalian orders, some families exhibited rapid protein evolution at better conserved positions across a substantial number of tree edges, while the opposite held true for other families. Here, Bradypodidae (three-toed sloths), Cercopithecidae (Old World monkeys), and Octodontidae (encompassing degus, rock rats, and viscacha rats) were associated with the three highest median θ_evo_ values (Figure 3A), while Equidae (horses and related animals), Leporidae (rabbits and hares), and Cervidae (deer) exhibited the three lowest median θ_evo_ values. Calculation of 90% confidence intervals supported the ranking of families obtained by calculation of median θ_evo_ values (Figure 3B and 3C). Analysis of individual OXPHOS complexes provided similar results regarding the relative magnitudes of mitochondria protein evolution among mammalian families (Supplementary Figs. S4, S5, and S6).

**Figure 3:**
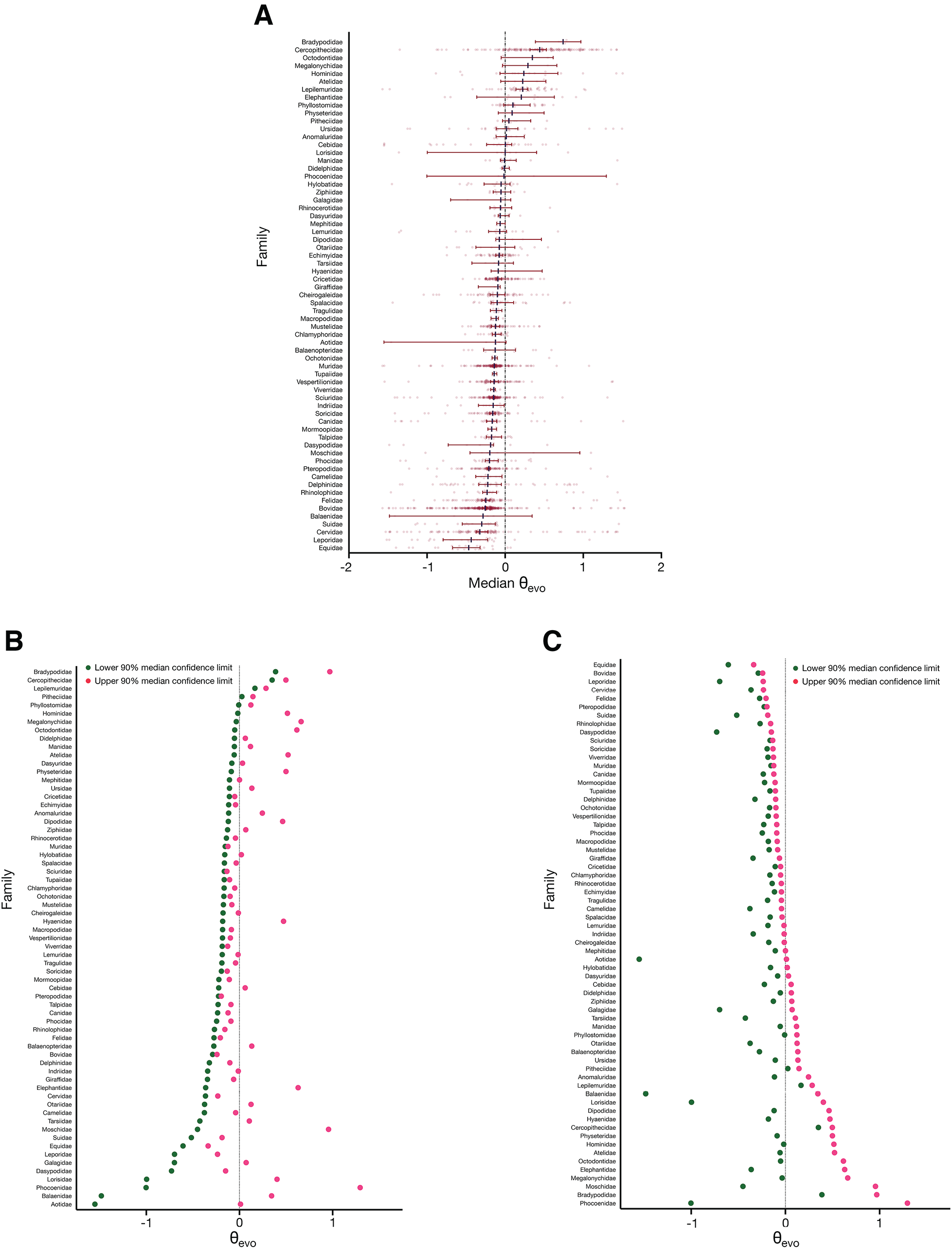
Mammalian families differ in their likelihood of potentially consequential mitochondrial protein substitutions. The analyses in (**A**), (**B**), and (**C**) were performed as in Figure 2, except mammalian families assigned five or more edges are analyzed.

Given the abundance of mitochondrial protein evolution found among primates, we focused additional attention upon this particular clade by examining the trends among primate suborders, infraorders, subfamilies, and genera. At the rank of suborder, Haplorrhini (which includes Old World monkeys, New World monkeys, and apes) manifested a higher median θ_evo_ value than Strepsirrhini (encompassing lemurs, lorises, and galagos) (Figure 4A), a result supported by examining the 90% confidence limits of the median (Figure 4B). At the level of infraorder, Simiiformes (Old World monkeys, New World monkeys, and apes) exhibited a substantially higher median θ_evo_ value than other groups (Figure 4A), and this finding was also supported by examination of 90% confidence intervals (Figure 4B). When considering subfamilies, Cercopithecinae (encompassing most Old World monkeys, including baboons, macaques, and vervet monkeys) provided the clearest evidence for abundant and likely consequential mitochondrial protein evolution, although the θ_evo_ values within several other subfamilies, such as Cebinae (New World capuchin monkeys) and Colobinae (multiple genera of Old World monkeys) were also elevated (Figure 4C and Figure 4D). Finally, when genera were analyzed, substantive mitochondrial protein change was clearly evident within *Macaca* (macaques) (Figure 4E and Figure 4F). Median θ_evo_ values were also elevated for the genera *Chlorocebus* (encompassing vervet and green monkeys), *Gorilla* (gorillas), *Papio* (baboons), and *Trachypithecus* (lutungs).

**Figure 4:**
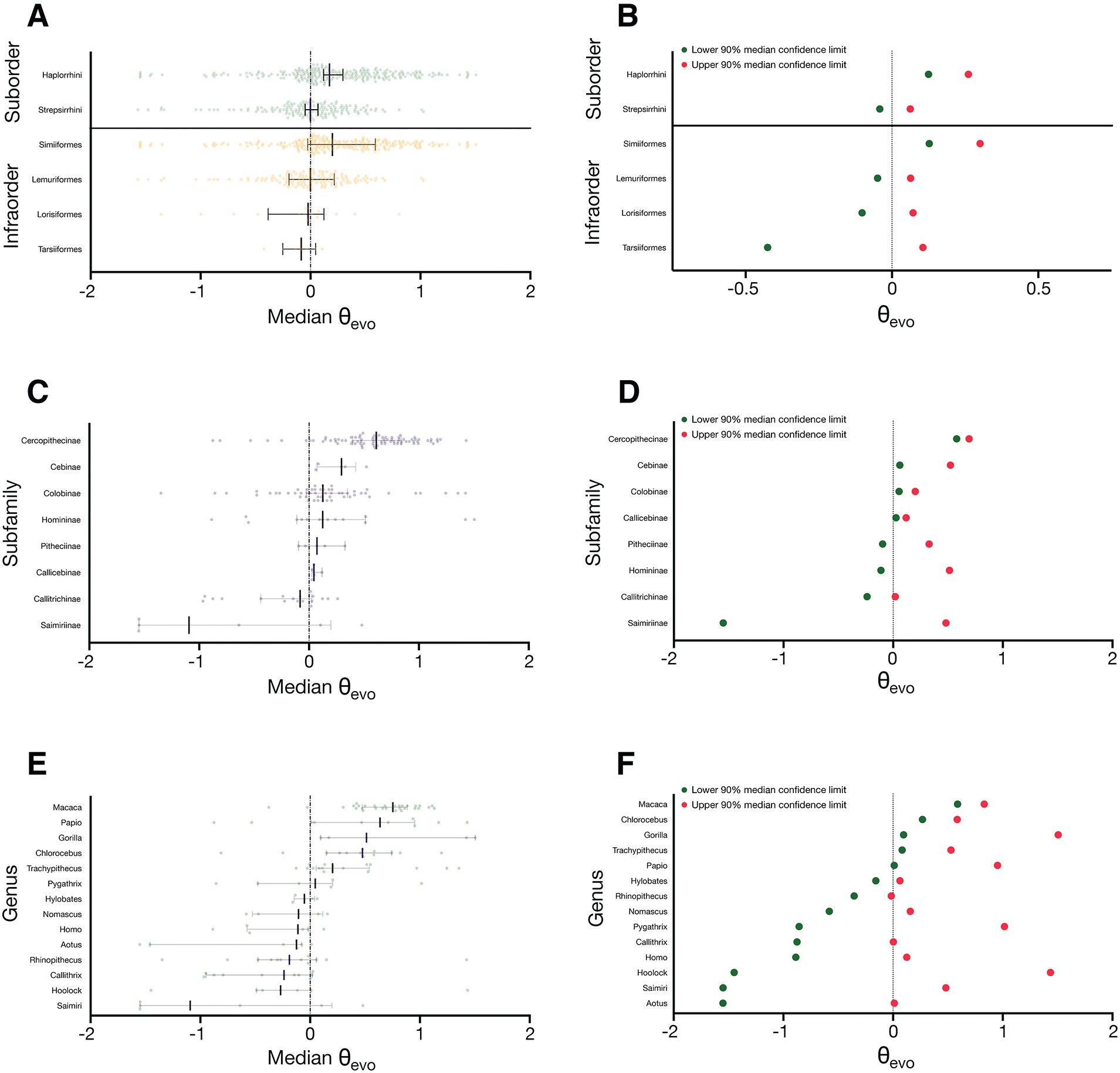
Substantial mitochondrial protein evolution is prevalent across several primate clades. Analyses were carried out as in Figure 2 and Figure 3, except primate suborders and infraorders (**A** and **B**), subfamilies (**C** and **D**), or genera (**E** and **F**) assigned five or more edges are included.

We hoped to better understand those substitutions (a particular ancestral character changed to a particular descendent character) that we identified along primate tree edges at the most conserved positions. Therefore, we tabulated all primate substitutions, then found matching substitutions within other mammalian orders. As successive TSS cut-offs associated with increasing site conservation were imposed upon this list of primate substitutions, the enrichment of the remaining substitutions within other orders relative to a list not subjected to TSS filtering was calculated. Intriguingly, as the TSS cut-off applied to primate substitutions was decreased toward a value of five or lower, an enrichment within specific clades became very clear. Concretely, the five orders Hyracoidea (hyraxes), Proboscidea, Microbiotheria (the monito del monte), Monotremata (includes platypuses and spiny anteaters), and Pilosa were most enriched with primate substitutions occurring at the most conserved positions (Figure 5A). When focusing our analysis upon substitutions at highly conserved sites found within haplorhines (Figure 5B) and anthropoids (Figure 5C), the results were quite similar, with Diprotodontia (which includes multiple marsupial families) replacing Proboscidea among the five top-ranked orders.

**Figure 5:**
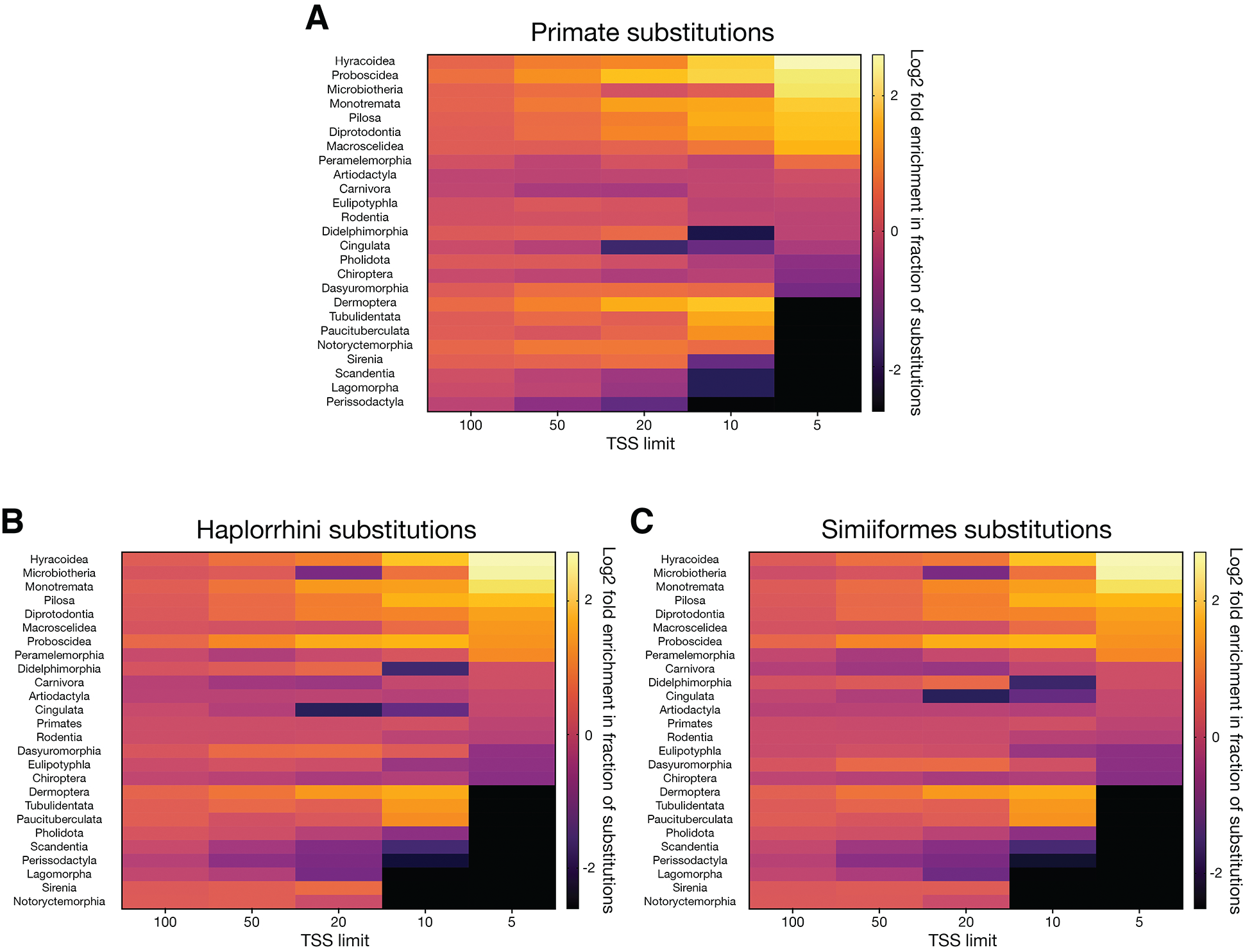
A search for specific primate substitutions at highly conserved mitochondrial protein positions outside of selected clades. Substitutions found along edges assigned to (**A**) primates, (**B**) haplorhines, and (**C**) anthropoids were tabulated, and those substitutions with the same ancestral and descendant characters were located among the various mammalian orders. The indicated TSS cut-offs were applied to this set of substitutions. The log2 fold change in the fraction of selected substitutions found in each order following the application of each indicated TSS cut-off, as compared to a dataset unfiltered by TSS, is indicated in the heat map.

Since anthropoids were replete with substitutions at positions highly conserved across mammals, we looked closely at the substitutions inferred along the most ancestral anthropoid tree edges at amino acid positions of TSS five or below, since the identification and characterization of those ancestral mutations should allow speculation regarding the metabolism and lifestyle of early anthropoids and their descendants. When focusing upon the branch leading to Old World monkeys, apes, and New World monkeys (edge **i**, Figure 6), a number of changes to highly conserved positions with a TSS of five or less were inferred. Intriguingly, all of these mutations were found within Complex IV, consistent with previous analyses highlighting substantial change to primate cytochrome *c* oxidase components (Gissi et al. 2000; Grossman et al. 2004; Pierron et al. 2011, 2012). Additional changes likely to be functionally relevant, mostly located within Complex IV, were found along the descendent edges leading toward Old World monkeys and apes (edge **ii**, Figure 6), or toward New World monkeys (edge **iii**, Figure 6).

**Figure 6:**
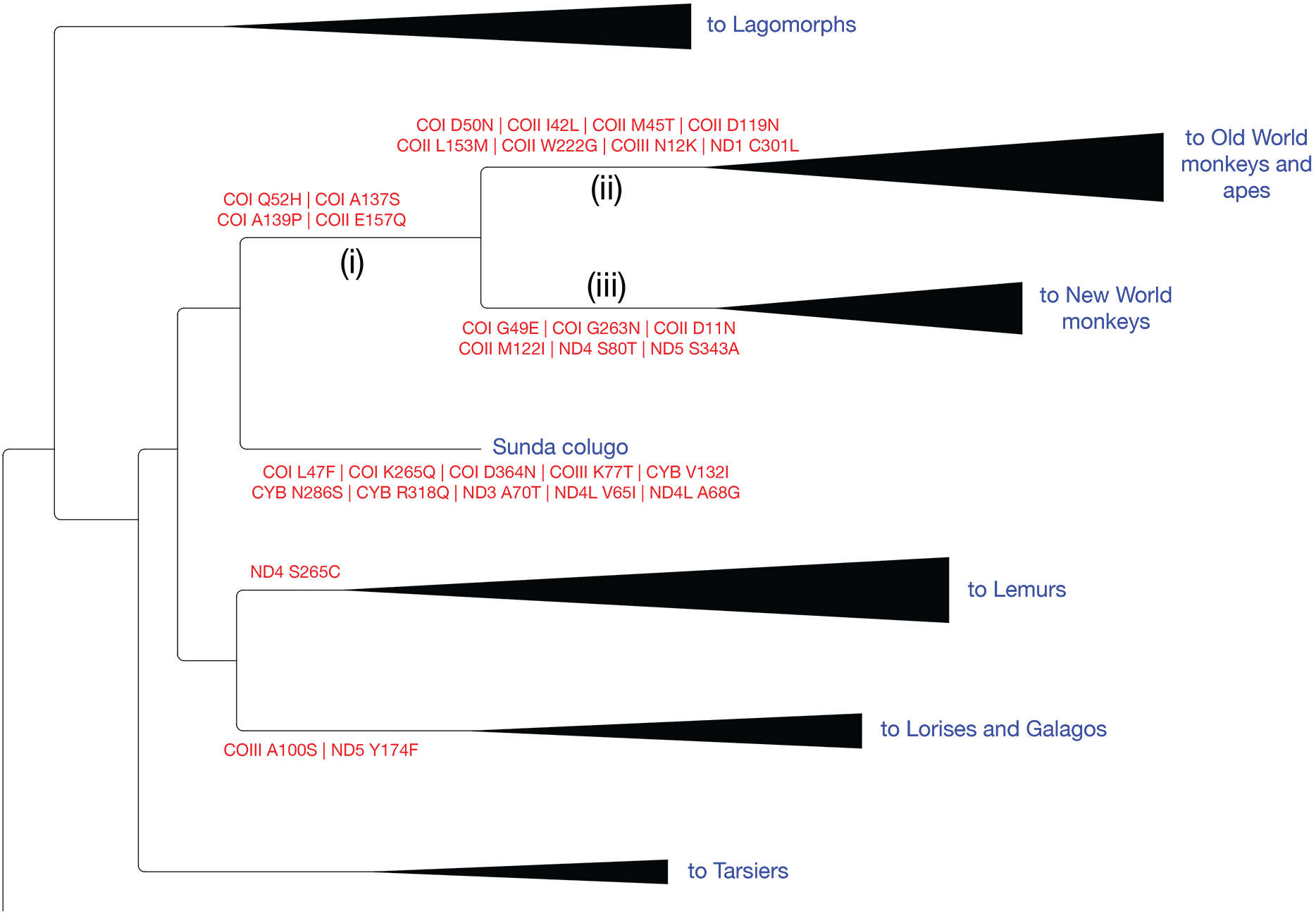
Substitutions at highly conserved mitochondrial protein positions are predicted at edges representing early anthropoid evolution. A portion of the inferred mammalian mtDNA tree is provided, and all substitutions occurring at positions with a TSS of five or lower are shown. Edge (**i**) leads to the node parental to Old World monkeys, apes, and New World monkeys, edge (**ii**) leads to Old World monkeys and apes, and edge (**iii**) leads to New World monkeys. All alignment positions are indexed to *Bos taurus* reference sequences.

We then sought mutations outside of primates that would precisely match those substitutions, by both ancestral and descendant amino acid, inferred along these three ancestral anthropoid edges at highly conserved (TSS ≤ 5) positions. Along the edge leading to Old World monkeys, apes, and New World monkeys (edge **i**, Figure 6), a Q52H change in *Antilope cervicapra* (blackbuck), a species characterized by a very high running speed, was predicted (Table 1). However, when considering the descendent edges leading to Old World monkeys and apes (edge **ii**, Figure 6) or to New World monkeys (edge **iii**, Figure 6), most of the substitutions at highly conserved sites matched those found within, or in edges leading to, groups quite conspicuous for their reduced basal metabolism, such as tuco-tucos, koalas, naked mole-rats, three-toed sloths, vampire bats, anteaters, and the monito del monte (Table 1).

**Table 1:** Substitutions found outside of primates that match those in selected primate ancestral edges.

| Edge analyzed | Edge description | Complex | Substitution (*Bos taurus* indexed) | Matching edge | Comment |
| --- | --- | --- | --- | --- | --- |
| i | Edge leading to Old World monkeys, apes, and New World monkeys | IV | COX1 Q52H | to *Antilope cervicapra* | *Antilope cervicapra* has a very high maximal running speed (Lovegrove 2004). |
| ii | Edge leading to Old World monkeys and apes | I | ND1 C301L | Myrmecophagidae internal edge | Daughter edges lead to *Tamandua tetradactyla*, *Tamandua mexicana*, *Myrmecophaga tridactyla*. Anteaters typically exhibit a reduced basal metabolic rate and body temperature relative to expectation (Stahl et al. 2012; McNab 2009). |
| ii | Edge leading to Old World monkeys and apes | IV | COX2 I42L | Dasyuromorphia internal edge | Daughter edges encompass a number of marsupial families. Marsupials typically have basal metabolic rates that are lower than placental mammals. Several species descending from this edge undergo torpor, a low-metabolism, low-temperature, decreased-activity state. (MacMillen and Nelson, 1969; Geiser 1994). |
| ii | Edge leading to Old World monkeys and apes | IV | COX2 I42L | to *Dromiciops gliroides* | *Dromiciops gliroides* is a marsupial with a basal metabolic rate below that predicted for a placental mammal of similar size. This organism can enter a prolonged torpor state induced by low ambient temperature and reduced food intake (Bozinovic et al. 2004). |
| ii | Edge leading to Old World monkeys and apes | IV | COX2 I42L | to *Murina ussuriensis* | *Murina ussuriensis* is the only bat known to hibernate in snowbanks, a location may provide greater temperature stability during hibernation (Hirakawa and Nagasaka 2018). |
| ii | Edge leading to Old World monkeys and apes | IV | COX2 M45T | *Bradypus* internal edge | Members of the *Bradypus* genus manifest greatly reduced basal metabolic rates and low activity relative to other mammals, and the body temperatures of these sloths can fluctuate wildly (Cliffe et al. 2018; Pauli et al. 2016; Britton and Atkinson 1938). |
| ii | Edge leading to Old World monkeys and apes | IV | COX2 M45T | to *Elephas antiquus* | *Elephas antiquus* (or *Palaeoloxodon antiquus*) is an extinct elephant of greater size than extant elephants. Larger mammals tend to have lower basal metabolic rates than smaller mammals, and elephants have a lower energy expenditure and basal metabolic rate than its size would predict (McNab 2012; Christiansen 2004) |
| iii | Edge leading to New World monkeys | I | ND4 S80T | Desmodontinae internal edge | Daughter edges lead to *Diphylla ecaudata*, *Diaemus youngi*, and *Desmodus rotundus*. Vampire bats typically have lower basal metabolism than other bats (McNab 2003). |
| iii | Edge leading to New World monkeys | I | ND5 S343A | Arvicolinae internal edge | Many arvicolids manifest a high basal metabolic rate and high body temperature, consistent with a high rate of energy expenditure (McNab 1992). |
| iii | Edge leading to New World monkeys | I | ND5 S343A | Bathyergidae internal edge | Daughter edges lead to the fossorial rodents *Heterocephalus glaber* (naked mole-rat) and *Fukomys damarensis* (Damaraland mole-rat). *Heterocephalus glaber* exhibits a basal metabolic rate far lower than expectations, is a poor thermoregulator, and is highly tolerant of hypoxia (McNab 1966; Park et al. 2017). |
| iii | Edge leading to New World monkeys | IV | COX1 G49E | to *Phascolarctos cinereus* | The basal metabolic rate of koalas are only ~74% of expectations for marsupials of its size, and koalas are characterized for their low activity (Degabriele and Dawson 1979; Nagy and Martin 1985). |
| iii | Edge leading to New World monkeys | IV | COX1 G49E | to *Ctenomys leucodon* | Tuco-tucos are fossorial rodents. Typically, tuco-tucos exhibit lower-than-expected basal metabolic rates (Busch 1989). |

<sup>a</sup> Those substitutions matching mutations inferred at the ancestral edges (i), (ii), and (iii) shown in Fig. 6 are listed, along with comments regarding the metabolic proficiency of the associated clade or species.

## Discussion

We mapped predicted amino acid substitutions to the edges of a tree inferred from the coding regions of mammalian mitochondrial genomes. Using this substitution map, we calculated the TSS, a measure of conservation robust against sampling biases and benefitting from a high dynamic range, for alignment positions within mitochondria-encoded proteins. Using those TSSs, our substitution map, and branch lengths determined from protein coding mtDNA sequences, we calculated a score (θ_evo_) that reports upon whether an unusual amount of functional change may have occurred along a phylogenetic tree edge. Specific substitutions along high-ranking tree edges that may be most impactful, based upon their location at conserved positions, are easily retrieved for further analysis. To our knowledge, we have carried out the most comprehensive analysis of mammalian mtDNA-encoded protein sequences performed to date, and our findings serve as a useful resource for those investigating the structure and function of mitochondria-encoded OXPHOS components. In addition, our dataset is highly valuable for those researchers interested in selective forces that may have helped shape mtDNAs during the establishment and expansion of diverse mammalian clades.

Other approaches have previously been deployed toward the detection of substitutions linked to functional change within mitochondrial proteins (Garvin et al. 2015). For example, the dN/dS ratio, a comparison of non-synonymous to synonymous nucleotide change, serves as the basis for several popular methods seeking unusually rapid evolution of proteins (Suzuki and Gojobori 1999; Yang et al. 2000; Kosakovsky Pond and Frost 2005). However, analysis of large datasets by this approach can be computationally intensive (Jeffares et al. 2015), and the ability of this metric to detect positive or relaxed selection on a protein may be diminished when many sites are highly conserved (Sharp 1997; Hughes 2007; Echave et al. 2016). Other weaknesses further limit the utility of dN/dS-based approaches (Kryazhimskiy and Plotkin 2008; Spielman and Wilke 2015; Venkat et al. 2018; Wisotsky et al. 2020), including a difficulty in identifying instances of fixed adaptive substitutions subject to stabilizing selection, even though these changes are often linked to habitation of a new niche or to an interesting evolutionary transition (Hansen 1997; Bloom 2017). Alternative methodologies can take advantage of the chemical properties of amino acids to identify adaptive change in polypeptides (Woolley et al. 2003; McClellan et al. 2005). However, generalized measures of amino acid side behavior may not always allow detection of functional change, since the site-specific acceptability of a given substitution is determined by its local protein environment (Zuckerkandl and Pauling 1965). Our rapid and computationally inexpensive approach will complement these established methods that seek notable change to mitochondrial polypeptides, and we certainly anticipate the application and further development of our methodology outside of the study of bioenergetics.

Orders and families can encompass members encountering a wide spectrum of ecological factors and life histories, and we indeed encounter many outliers when examining the θ_evo_ values found within each taxonomic group. However, as might be expected, based upon generations of selection for survival within divergent ecological niches, metabolic characteristics differ among taxonomic groups (McNab 2008; Griebeler and Werner 2016), and we have noted a tendency for athleticism and/or high basal metabolic rates to correspond with the conservation of mtDNA-encoded OXPHOS complex members among land mammals. Indeed, artiodactyls and lagomorphs exhibit the greatest residual basal metabolic rates among mammals (Hayssen and Lacy 1985; Lovegrove 1996, 2000; McNab 2008; Garland 2009), and we detect the strongest conservation of mtDNA-encoded proteins among these orders. In contrast, many changes rapidly accumulate at conserved mitochondrial protein positions when considering tree edges assigned to the order Pilosa and family Bradypodidae. Accordingly, sloths are also characterized by very low basal metabolic rates and poor thermoregulation (Britton and Atkinson 1938; Pauli et al. 2016; Cliffe et al. 2018). Substantial change to positions conserved among mammals is also detected across primates, and we discuss the evolution of primate mitochondrial proteins below. Cladistic differences in the substitution profiles of mitochondrial proteins support the idea that metabolism can be subject to phylogenetic inertia that may hamper adaptation to new diets and climes (Westoby et al. 1995; Hansen 1997).

### Primates are enriched for mitochondrial protein substitutions that are predicted to have functional effects

Consistent with the results described above, previous studies of primates, necessarily more limited in scale due to contemporary limitations in sequence availability, have also revealed accelerated evolution and unusual substitutions within mtDNA- and nucleus-encoded OXPHOS complex subunits (Gissi et al. 2000; Pupko and Galtier 2002; Grossman et al. 2004; Pierron et al. 2011, 2012). At lower taxonomic levels, we found the clearest evidence of accelerated change to mtDNA-encoded proteins within the suborder Haplorrhini, the infraorder Simiiformes, and several families, subfamilies, and genera spread distributed throughout the primate tree. When those substitutions occurring at the most conserved positions within primate, haplorhine, or anthropoid tree edges were sought outside of these groups, there was a substantial enrichment within the orders Hyracoidea, Proboscidea, Microbiotheria, Monotremata, Pilosa, and Diprotodontia. Low basal metabolism and difficulties in thermoregulation appear to typify the order Hyracoidea (Rübsamen et al. 1982). The related order, Proboscidea, includes elephants and their close relatives, which are also characterized by reduced basal and field metabolism relative to other mammals (Christiansen 2004). The order Microbiotheria contains one extant marsupial species, the monito del monte, which presents a low basal metabolic rate and can enter a prolonged torpor state in order to conserve energy (Bozinovic et al. 2004). The basal metabolic rate of monotremes is considerably lower than most placental mammals (Nicol 2017), with monotremes displaying the lowest body temperature among groups of mammals (Clarke and O’Connor 2014). The enfeebled metabolism of Pilosa members has been described above, and several members of the marsupial order Diprotodontia, such as the koala bear (Degabriele and Dawson 1979; Nagy and Martin 1985) and the wombat (Barboza et al. 1993; Evans et al. 2003), are low-activity mammals exhibiting reduced basal metabolism.

### Substitutions to the mitochondrial proteins of ancestral anthropoids are likely to compromise OXPHOS activity

Next, we looked specifically at mutations occurring along three selected edges representing the origin of anthropoids. The majority of those mutations at the most conserved positions were found in Complex IV, and a subset of these were previously placed at the interaction surface between cytochrome *c* and cytochrome *c* oxidase (Schmidt et al. 2005). Complex IV is a key control point for OXPHOS activity (Piccoli et al. 2006; Dalmonte et al. 2009; Arnold 2012; Hill 2016), and given the scarcity of these selected mutations among other clades, compensation allowing functional neutrality (Starr and Thornton 2016) would, in our view, be highly unlikely. Instead, alteration of Complex IV activity by these substitutions is a more plausible scenario. For example, a change similar to the ancestral anthropoid substitution COII D119N, where a negatively charged side chain is replaced by a neutral side chain, reduces cytochrome *c* oxidase activity in the alpha-proteobacterium *Rhodobacter sphaeroides* (Wang et al. 1999; Zhen et al. 1999), and there is no clear evidence that this change might be compensated in the apposed, nucleus-encoded cytochrome *c* protein (Schmidt et al. 2005). At COI, the D50N (ancestral to Old World monkeys and apes) and G49E (ancestral to New World monkeys) substitutions are found near residue D51, which is subject to redox-dependent conformational and protonation changes (Yoshikawa et al. 1998; Okuno et al. 2003; Maréchal et al. 2012; Sharma et al. 2017) and forms a key component of the ‘H-channel’ that serves as either a proton transfer pathway or as a dielectric well (Yoshikawa et al. 1998; Rich and Maréchal 2013; Sharma et al. 2017). The Q52H (ancestral to anthropoids) substitution adds a titratable histidine to a region important for proton and water release (Son et al. 2017; Cai et al. 2018). All three of these COI substitutions, quite unusual among mammals, should be expected to alter enzyme kinetics.

If those mutations in stem anthropoids did lead to changes in OXPHOS activity, might they have boosted or reduced cellular metabolism? Equivalent substitutions to those found along these three edges, in findings that mirror those obtained by our broader analysis of all primate, haplorhine, or anthropoid edges, were found among clades dominated by species with poor temperature regulation and/or low basal metabolic rates relative to the bulk of mammals (Britton and Atkinson 1938; McNab 1966, 2003, 2009, 2012; Degabriele and Dawson 1979; Nagy and Martin 1985; Busch 1989; Christiansen 2004; Bozinovic et al. 2004; Stahl et al. 2012; Pauli et al. 2016; Cliffe et al. 2018). Specifically, inferred substitutions found along the tree edge leading to Old World monkeys could be matched in Myrmecophagidae (anteaters), *Dromiciops* (monito del montes), *Bradypus* (three-toed sloths), and *Elephas* (elephants). Inferred mutations found along the tree edge leading to New World monkeys were found in Desmodontinae (vampire bats), Bathyergidae (mole rats), *Phascolarctos* (koalas), and Ctenomys (tuco-tucos). Taken together, our analysis of mammalian mtDNAs suggests a general reduction in the activity of OXPHOS complexes encoded by ancestral, and perhaps a substantial number of extant, anthropoids. In contrast, evidence that the changes to mtDNA-encoded proteins among primates would generally lead an increase in OXPHOS activity is scant, although it is certainly possible that mtDNA changes in some lineages, including reversion of more ancestral substitutions that had previously limited metabolic activity, may bypass the effects of more ancestral mutations that limit OXPHOS component activity. Indeed, many anthropoids exhibit basal metabolic rates that comport with general expectations for mammals (Pontzer et al. 2014; Rottenberg 2014).

If the mtDNA-encoded proteins of early anthropoids were characterized by lower OXPHOS complex activity and reduced cellular metabolism, how might the expansion of an energetically expensive (Mink et al. 1981; Harris et al. 2012; Bonvento and Bolaños 2021) primate brain be rationalized?

The basal metabolic rates of whole organisms cannot be *directly* linked to encephalization (Roth and Dicke 2005), or the extent to which the brain size corresponds to expectations based upon size (Jerison 2012). For example, monotreme encephalization generally appears to fall in line with that of placental mammals (Macrini et al. 2006; Ashwell and Ashwell 2013; Nicol 2017), in spite of their comparatively reduced metabolism. Elephants also exhibit lower-than-expected basal metabolic rates (Christiansen 2004), yet elephants are more highly encephalized than many mammals (Hofman 1982; Roth and Dicke 2005). Lagomorphs, which exhibit a relatively high basal metabolic rate (Hayssen and Lacy 1985; Lovegrove 2000) tend not to be highly encephalized (Kruska 2005; Roth and Dicke 2005).

If brain expansion is not always coincident with high basal metabolism, then one might speculate instead (Pontzer 2015), consistent with the expensive tissue hypothesis (Aiello and Wheeler 1995; Aiello et al. 2001), that cellular physiology and OXPHOS components were initially selected for energetic parsimony due to limited food availability. Low expenditure of energy by cellular metabolism then licensed later evolution of a larger brain upon a subsequent, sustained improvement in diet. In this case, the addition of new brain tissue prompted a return to mass-specific metabolic rates across the whole organism that fall within general expectations for mammals. Further support for this idea will certainly require additional comparison of the mitochondrial activities and the metabolic properties of primates with those of other mammalian groups.

## Materials and Methods

### Sequence acquisition and concatenation

Record accessions for mammalian reference mtDNAs were obtained using the Organelle Genome Resources provided by the National Center for Biotechnology Information Reference Sequence project (NCBI RefSeq, Release 204, https://www.ncbi.nlm.nih.gov/genome/organelle/) (O’Leary et al. 2016). All records not containing ‘NC_’ at the beginning of their accession name were removed, then the accession for the reptile *Anolis punctatus* was added. Using this list of accessions, full GenBank records for these mtDNAs were downloaded by use of the NCBI Batch Entrez server (https://www.ncbi.nlm.nih.gov/sites/batchentrez). This file (Supplementary File 4) was used as input for the ‘vert_mtDNA_genbank_grab.py’ script (https://github.com/corydunnlab/mammal_mitoprot_evolve).

### Sequence alignment

The amino acid sequences of individual mitochondrial proteins extracted from the GenBank input file, as well as a concatenate of protein coding sequences, was aligned using the FFT-NS-2 method in MAFFT v7.475 (Katoh and Standley 2013).

### Phylogenetic tree inference

Our alignment of mtDNA sequences encoding mitochondrial proteins was used as input for RAxML-NG v0.9.0 (Kozlov et al. 2019) under a single partition, a GTR+FO+G4m model of DNA change, and a seed set as 200. 10 random and 10 parsimony-based starting trees were used to initiate tree inference, and the average relative Robinson-Foulds distance (Robinson and Foulds 1981) for the resulting set of trees was 0.01. The best scoring tree is provided as Supplementary File 5. 600 bootstrap replicates were generated using RAxML-NG, and a weighted Robinson-Foulds distance converged below the 1% cutoff value. Both Felsenstein’s Bootstrap Proportions (Supplementary File 6) (Felsenstein 1985) and, perhaps more appropriate for our large dataset (Soltis and Soltis 2003), the Transfer Bootstrap Expectations (Supplementary File 7) (Lemoine et al. 2018) were calculated and used to label our best scoring tree. The best maximum likelihood tree was used for downstream analyses after using FigTree 1.4.4 (https://github.com/rambaut/figtree/releases) to place the root upon the branch leading to *Anolis punctatus*. We note that an inferred tree of mtDNA sequences should be expected to differ from other trees generated using nuclear DNA sequences subject to recombination or trees incorporating morphological characters.

### Calculation and classification of edge θ_evo_ values

GenBank records, sequence alignments of mitochondrial proteins, and our best maximum likelihood tree were used as input for the script ‘mtDNA_protein_evolution_calculator.py’ (https://github.com/corydunnlab/mammal_mitoprot_evolve). Within this script, PAGAN v0.61(Löytynoja et al. 2012) is called to perform ancestral prediction of protein sequences at internal tree nodes, and the resulting PAGAN alignments were ungapped using the *Bos taurus* reference sequence (NC_006853.1). A PAGAN output tree is provided as Supplementary File 8. Taxonomy of input accessions was recovered from the NCBI Taxonomy Database (access date: January 29, 2021). Taxonomy of internal edges at a given taxonomic level was determined by unanimous agreement among descendent edges. If multiple taxa were found downstream of a descendent node, ‘MIXED’ was assigned to that edge at the analyzed taxonomic level. Edge θ_evo_ values were further examined and plotted in GraphPad Prism v9.0.2, and only taxonomic groups containing five or more edges (allowing calculation of confidence intervals of greater than or equal to 90%) are reported in our median-based analyses of specific taxonomic groups.

### Enrichment of specific substitutions from a chosen clade within other clades, assessed at different levels of conservation

Specific substitutions found within the order Primates, the suborder Haplorrhini, and the infraorder Simiiformes were extracted from a table containing all amino acid substitutions at analyzed positions and their associated taxonomic assignments (Supplementary File 1). These same substitutions were then sought among edges found outside of these selected groups. The recovered dataset was filtered by TSS to determine the enrichment of anthropoid substitutions in other clades at different levels of conservation. These analyses were performed using the scripts ‘primates_specific_substitutions_outside_of_primates_with_TSS_enrichment.py’, ‘haplorrhini_specific_substitutions_outside_of_haplorrhini_with_TSS_enrichment.py’, and ‘simiiformes_specific_substitutions_outside_of_simiiformes_with_TSS_enrichment.py’ (https://github.com/corydunnlab/mammal_mitoprot_evolve).

### Detection of specific substitutions extracted from a selected edge across the remainder of a phylogenetic tree

We identified substitutions along the selected tree edge at alignment positions with TSSs below the selected limit, then matched those substitutions with those found at other locations in the tree. To complete this task, we provided input consisting of a table containing all amino acid substitutions at analyzed positions and their associated taxonomic data (Supplementary File 1), the selected reference edge, and a desired TSS threshold to the script ‘TSS_five_or_below_inside_and_outside_selected_edge.py’ (https://github.com/corydunnlab/mammal_mitoprot_evolve).

## Supporting information

Supplemental figures and captions for supplemental files

## Author contributions

C.D.D. conceived of the methodology. C.D.D. and B.A.A. wrote software. C.D.D. and V.S. analyzed data. B.A.A., V.S., and C.D.D. wrote the manuscript.

## Data availability statement

Supplementary figures and supplementary files harboring data underlying this article are available in Zenodo at https://doi.org/10.5281/zenodo.7265070. Scripts used for data analysis are found in GitHub in the repository found at https://github.com/corydunnlab/mammal_mitoprot_evolve.

## Acknowledgements

Funding for this project was obtained from the Sigrid Jusélius Foundation (Senior Researcher and Project Grants to C.D.D and V.S.), the Jane and Aatos Erkko Foundation (to C.D.D. and V.S.), the European Research Council (ERC Starting Grant RevMito 637649 to C.D.D.), the Academy of Finland (to C.D.D. and V.S.), the University of Helsinki (to V.S.) and the Magnus Ehrnrooth Foundation (to V.S). C.D.D. and V.S. also thank the Center for Scientific Computing, Finland (Puhti and Mahti supercomputers), the Information Technology for Science Group, University of Helsinki (Ukko2 cluster) and PRACE (Marconi100 hosted at CINECA, Italy) for essential computational support. We thank Gülayşe ince Dunn, Svetlana Konovalova, and Tamara Somborac for comments on the manuscript.

