## Supplemental figures and captions for supplemental files for "A novel approach to the detection of unusual mitochondrial protein change suggests low basal metabolism of ancestral anthropoids"

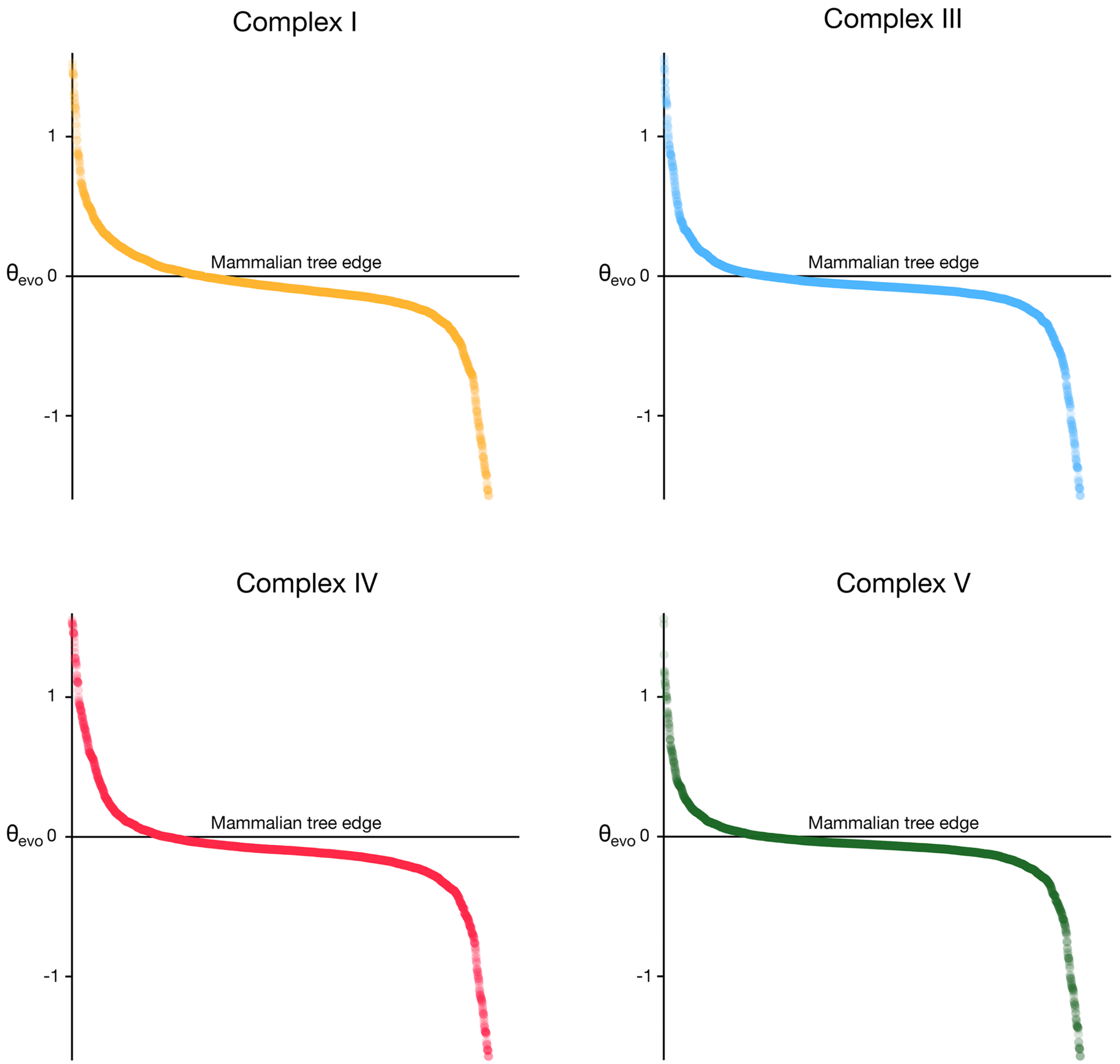


**Supplementary Figure S1: $\theta_{evo}$ calculated for each analyzed edge for specific OXPHOS complexes.** Analyses were performed as in Figure 1F, except that SPCSs calculated from mtDNA-encoded protein positions in Complex I, Complex III, Complex IV, or Complex V were used to generate $\theta_{evo}$ values.

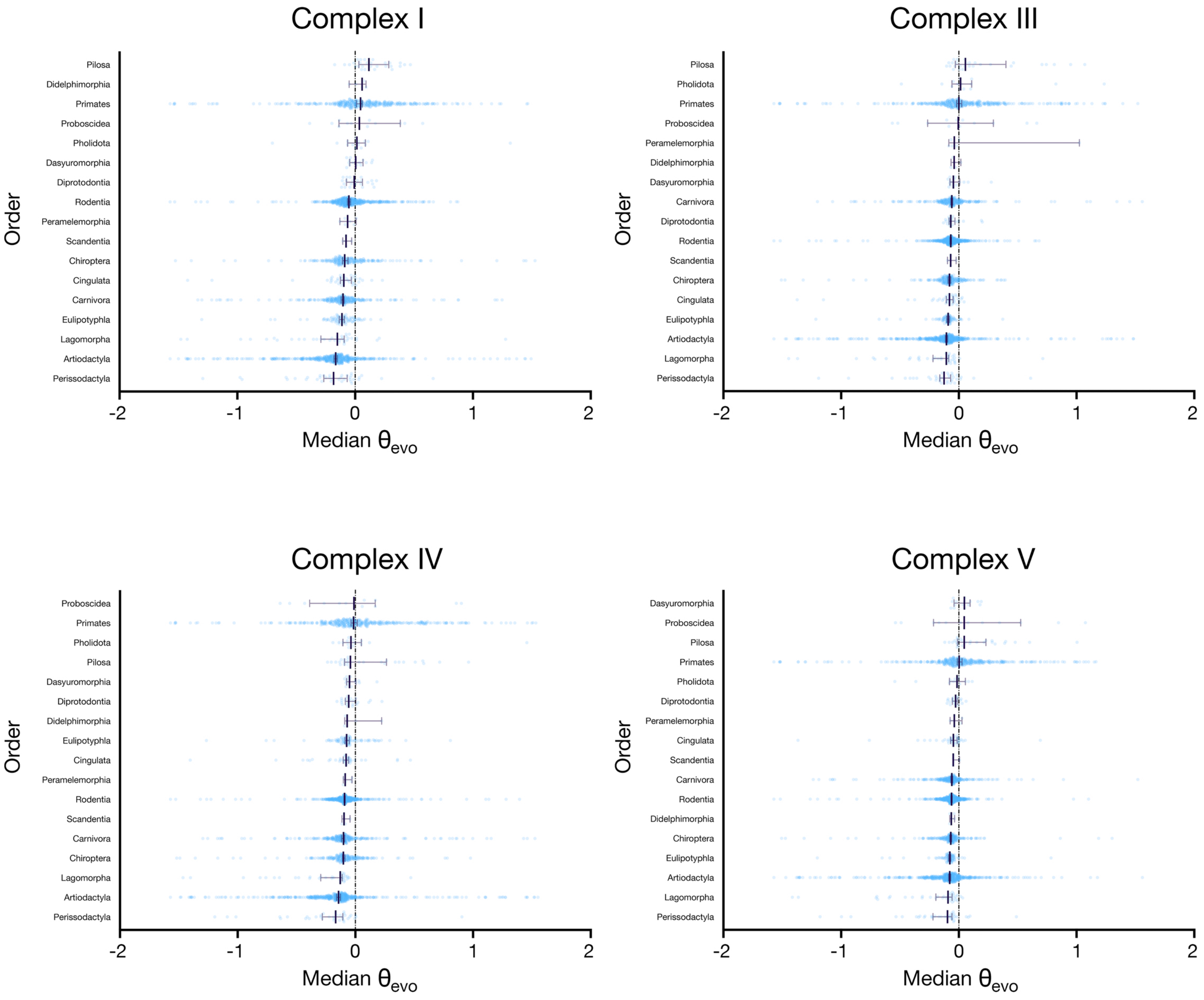


**Supplementary Figure S2: Mammalian orders differ in their propensity for potentially consequential mitochondrial protein substitutions within specific OXPHOS complexes (median calculations).** Analysis was performed as in Figure 2A, except that $\theta_{evo}$ values were obtained by analysis of mtDNA-encoded Complex I, Complex III, Complex IV, or Complex V polypeptides.

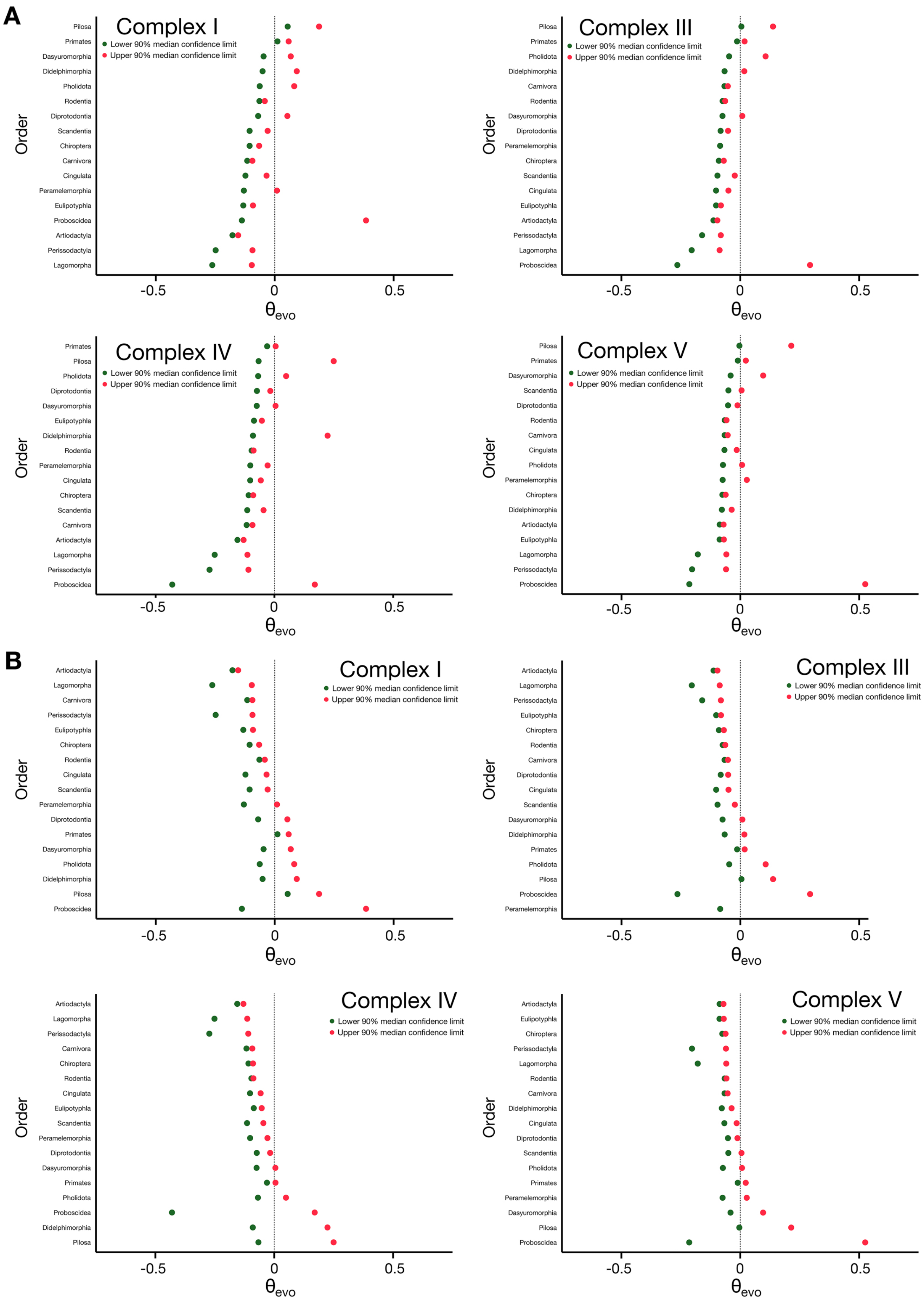


**Supplementary Figure S3: Mammalian orders differ in their propensity for potentially consequential mitochondrial protein substitutions within specific OXPHOS complexes (median confidence intervals).** Analysis was performed as in (**A**) Figure 2B or (**B**) Figure 2C, except that $\theta_{evo}$ values were obtained by analysis of mtDNA-encoded Complex I, Complex III, Complex IV, or Complex V proteins.

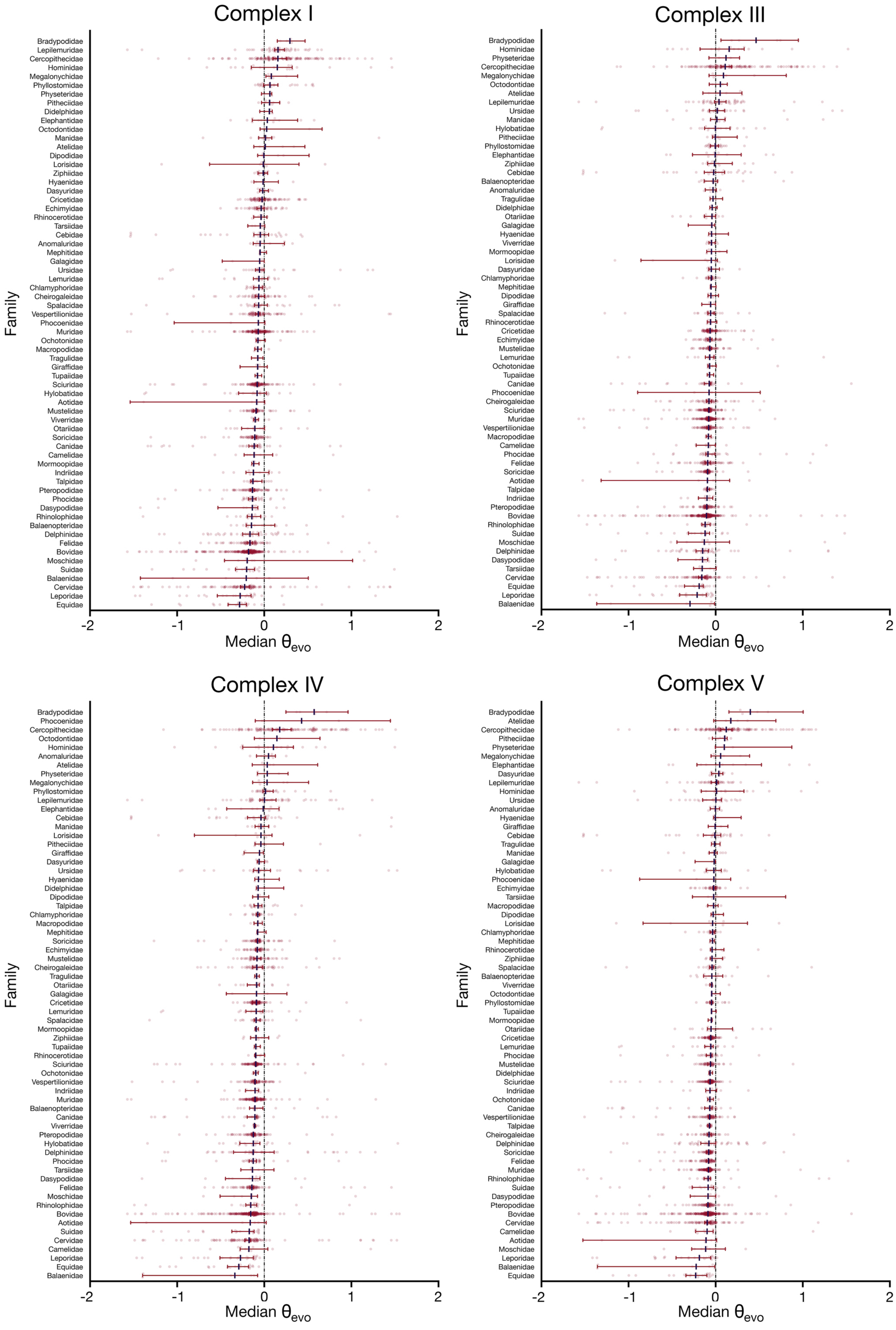


**Supplementary Figure S4: Mammalian families differ in their propensity for potentially consequential mitochondrial protein substitutions at specific OXPHOS complexes (median calculations).** Analysis was performed as in Figure 3A, except that $\theta_{evo}$ values were obtained by analysis of mtDNA-encoded Complex I, Complex III, Complex IV, or Complex V subunits.

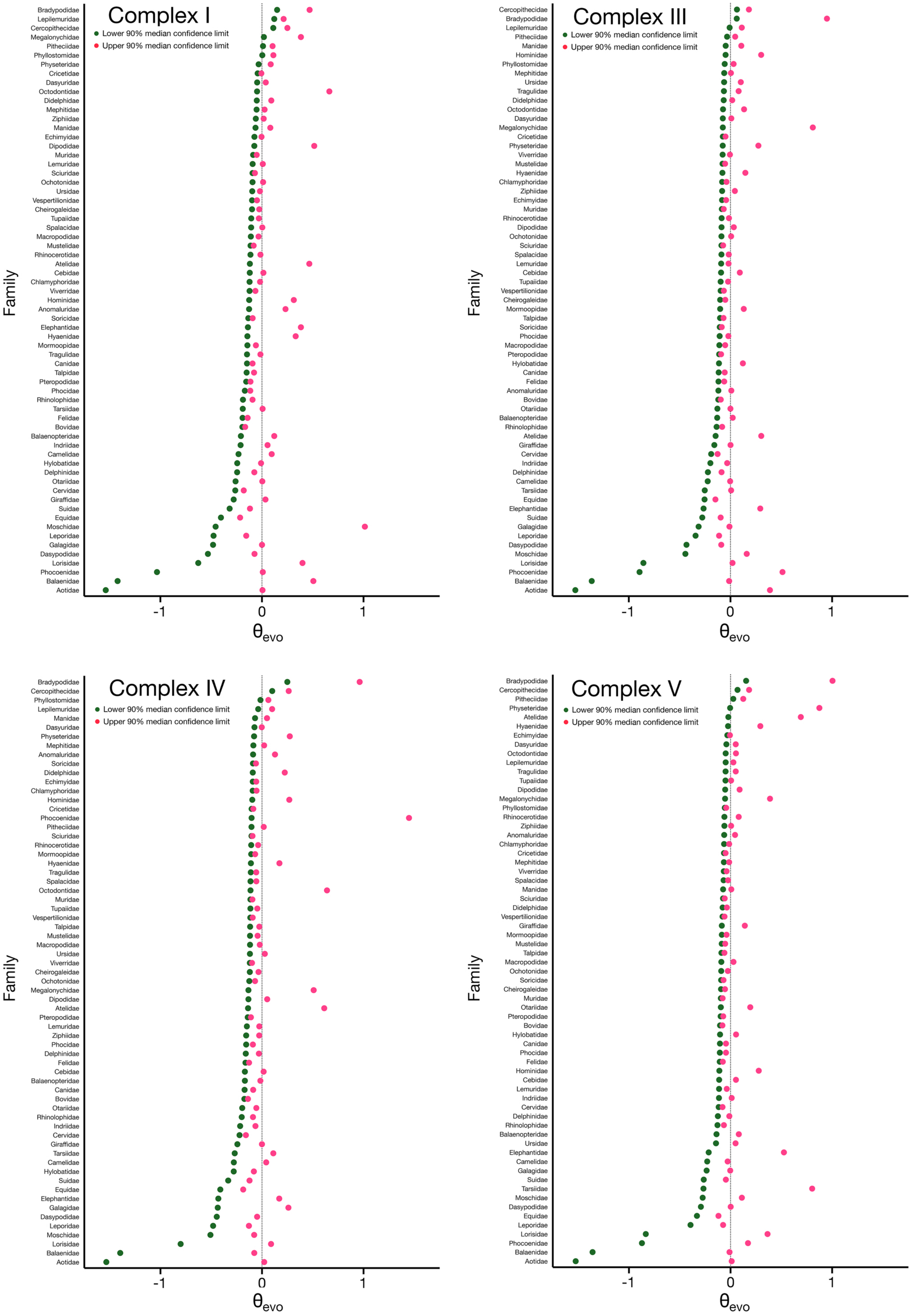


**Supplementary Figure S5: Mammalian families differ in their propensity for potentially consequential mitochondrial protein substitutions at specific OXPHOS complexes (median confidence intervals ordered by lower 90% median confidence limit).** Analysis was performed as in Figure 3B, except that $\theta_{evo}$ values were obtained by analysis of mtDNA-encoded Complex I, Complex III, Complex IV, or Complex V proteins.

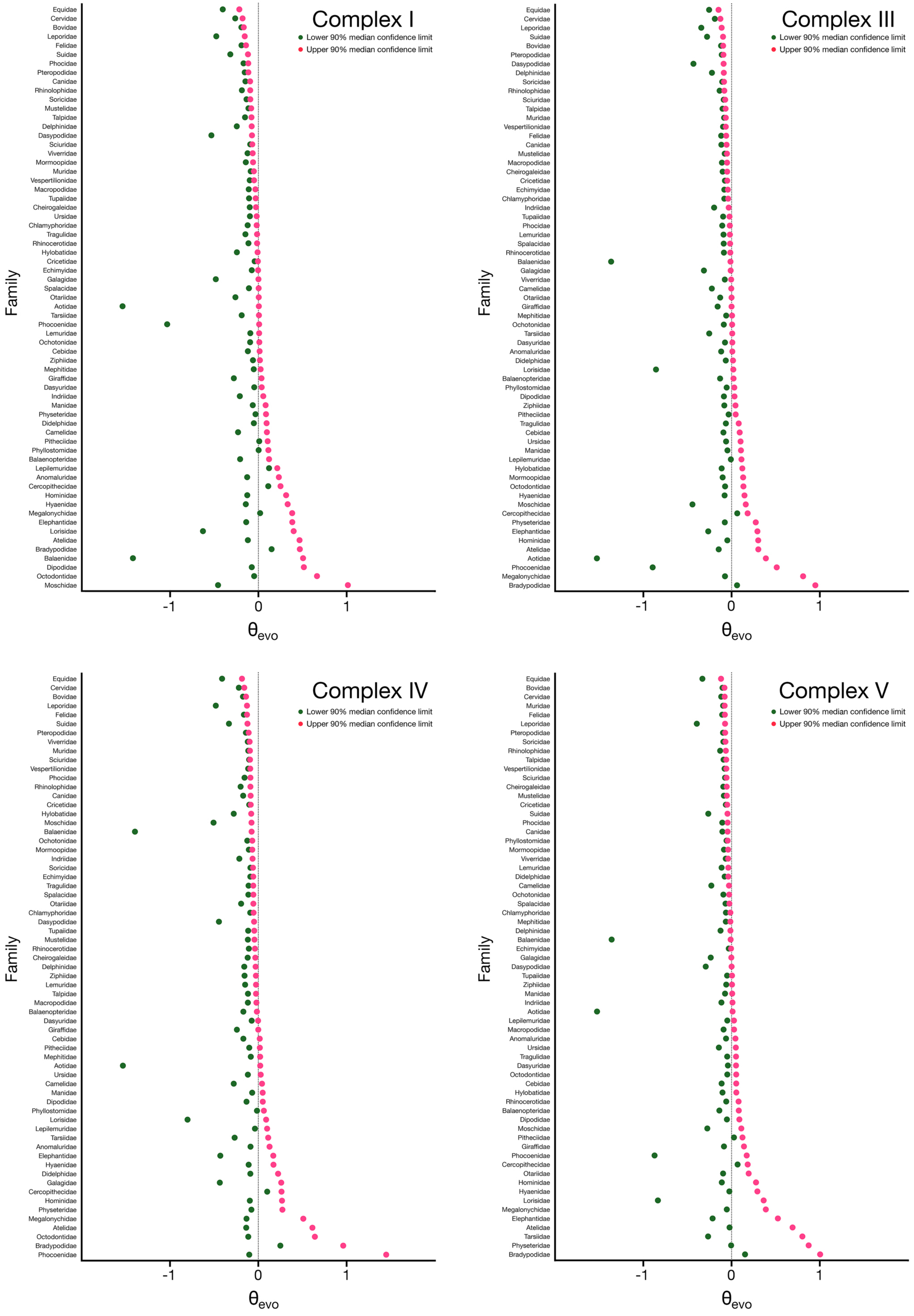


**Supplementary Figure S6: Mammalian families differ in their propensity for potentially consequential mitochondrial protein substitutions at specific OXPHOS complexes (median confidence intervals ordered by upper 90% median confidence limit).** Analysis was performed as in Figure 3C, except that $\theta_{evo}$ values were obtained by analysis of mtDNA-encoded Complex I, Complex III, Complex IV, or Complex V polypeptides.

**These supplementary files can be found at the following location:**

https://doi.org/10.5281/zenodo.7265070

**Supplementary File 1:** All predicted protein substitutions along all edges at positions containing less than 2% gaps across input and ancestral sequences are listed, along with associated taxonomy information, TSS, and branch length. All alignment positions refer to *Bos taurus* reference sequences.

**Supplementary File 2:** The TSS calculated for each mitochondrial protein alignment position. All alignment positions refer to *Bos taurus* reference sequences.

**Supplementary File 3:** SPCS and $\theta_{evo}$ outputs are provided for analyses across all mitochondria-encoded positions, as well as for focused analyses of specific OXPHOS complexes and individual proteins.

**Supplementary File 4:** A GenBank flat file containing RefSeq entries for mammalian mtDNAs, as well as the entry for the reptile *Anolis punctatus.*

**Supplementary File 5:** A maximum likelihood inferred tree generated by a RAxML-NG analysis of concatenated and aligned protein coding sequences from mammalian and *Anolis punctatus* mtDNAs.

**Supplementary File 6:** Bootstrap replicates were generated from the alignment of concatenated protein coding sequences. Felsenstein's Bootstrap Proportions (Felsenstein 1985) were calculated and used to label the maximum likelihood inferred tree of mammalian mtDNAs.

**Supplementary File 7:** Bootstrap replicates were generated using concatenated mammalian mtDNA coding sequences. Transfer Bootstrap Expectations (Lemoine et al. 2018) were calculated and used to label the maximum likelihood inferred tree of mammalian mtDNAs.

**Supplementary File 8:** PAGAN tree output produced using aligned amino acid sequences and the rooted maximum likelihood inferred tree as input.
